# Setting up an institutional OMERO environment for bioimage data: perspectives from both facility staff and users

**DOI:** 10.1101/2024.07.04.601832

**Authors:** Anett Jannasch, Silke Tulok, Chukwuebuka William Okafornta, Thomas Kugel, Michele Bortolomeazzi, Tom Boissonnet, Christian Schmidt, Andy Vogelsang, Claudia Dittfeld, Sems-Malte Tugtekin, Klaus Matschke, Leocadia Paliulis, Carola Thomas, Dirk Lindemann, Gunar Fabig, Thomas Müller-Reichert

## Abstract

Modern bioimaging core facilities at research institutions are essential for managing and maintaining high-end instruments, providing training and support for researchers in experimental design, image acquisition and data analysis. An important task for these facilities is the professional management of complex multi-dimensional bioimaging data, which are often produced in large quantity and very different file formats. This article details the process that led to successfully implementing the OME Remote Objects system (OMERO) for bioimage-specific research data management (RDM) at the Core Facility Cellular Imaging (CFCI) at the Technische Universität Dresden (TU Dresden). Ensuring compliance with the FAIR (findable, accessible, interoperable, reusable) principles, we outline here the challenges that we faced in adapting data handling and storage to a new RDM system. These challenges included the introduction of a standardized group-specific naming convention, metadata curation with tagging and Key-Value pairs, and integration of existing image processing workflows. By sharing our experiences, this article aims to provide insights and recommendations for both individual researchers and educational institutions intending to implement OMERO as a management system for bioimaging data. We showcase how tailored decisions and structured approaches lead to successful outcomes in RDM practices.

## Introduction

Bioimaging core facilities at universities and research institutions serve the scientific community by managing the acquisition, implementation, accessibility and reliability of modern high-end imaging instruments that are used by researchers on a daily basis. In addition, core facility staff provides training and support in using bioimaging instrumentation and analyzing the obtained image data, thereby contributing to numerous individual research projects. Beyond this, core facility staff is confronted with challenges concerning the professional management of bioimage research data.^1–3^ This data type, often derived from complex preparation protocols and imaging setups for delicate samples, is regarded as difficult to handle, store and share, because bioimage data are often multi-dimensional (time, space, channels, spectra, etc.). Moreover, bioimaging files are often large and acquired in proprietary file formats lacking a uniform data and metadata structure.^4^ Such challenges in data handling can be solved by investing in professional practices in bioimage research data management for core facilities. This needs to be achieved in close collaboration across institutional departments, including the central and/or departmental information technology (IT) units, institutional management boards and, of course, the user base of the facility.

The Core Facility Cellular Imaging (CFCI), a joint light and electron microscopy facility at the Faculty of Medicine Carl Gustav Carus at the Technische Universität Dresden (TU Dresden), is one of 12 core facilities around the local university hospital campus located in Dresden, Germany. These facilities collaborate through the multi-institutional bioimaging network, [Biopolis Dresden Imaging Platform](https://www.biodip.de/) (BioDIP). Due to sensitive patient-related data collected at the university hospital, the Faculty of Medicine has its own and independent IT department, and works in collaboration with the central IT department of the TU Dresden. To facilitate the adoption of bioimage data handling and management in compliance with the FAIR (findable, <u>a</u>ccessible, <u>i</u>nteroperable, <u>r</u>eusable) principles for CFCI users, we decided to pilot the local implementation of the bioimage-specific research data management system OME Remote Objects (OMERO). Currently, OMERO is the most widely used and best-established bioimage data platform.^5^ In line with our intention to implement new standards and routines for research data management (RDM) in our facility, we have contributed as a use case to two larger bioimaging RDM projects. First, within the [Information Infrastructure for BioImage Data](https://gerbi-gmb.de/i3dbio) (I3D:bio) project we have volunteered as a test site for the implementation of facility-oriented OMERO implementation guidelines. Second, as an early use case we collaborated with members of the NFDI4BIOIMAGE consortium, a part of Germany’s National Research Data Infrastructure (NFDI) to establish user-oriented support for metadata annotation. Based on the Recommended Metadata for Biological Images (REMBI), we connected image analysis workflows to OMERO. This was done in the interest of sharing publication-associated data in our OMERO instance.

In this article, we report in detail about our case in establishing OMERO as the basis for bioimaging research data management practices in our facility. As tailored decisions were required at multiple steps during the implementation process, our established single OMERO instance cannot be considered as a blueprint for all future implementations. However, we have leveraged a structured approach to implement OMERO as proposed by the I3D:bio project partners, and we report here on the reasoning and decision making processes that have culminated in the successful OMERO implementation at the CFCI of the Faculty of Medicine at the TU Dresden. Underlining this success, we further report on four different pilot-use cases at our faculty and how these users benefited from the tremendous possibilities offered by an integrated OMERO-based RDM approach.

## Initial institutional and administrative challenges

Since 2019 we have been engaged in research data management (RDM) with the objective to use OMERO as an open-source solution for handling, storing and sharing of the acquired image data.^5,6^ The latest technological developments in microscopy in line with the associated highly complex data structure and volumes require the establishment of RDM practices for bioimage data according to the FAIR principles^7^ before the data acquisition stage. Consequently, imaging facilities need to provide scientists with the appropriate tools to follow those practices. Through attending various lectures and workshops in the microscopy community and closely networking at regular meetings of the open exchange group, Research Data Management for Microscopy (RDM4mic), our facility staff became introduced and well acquainted with the functionality and benefits of OMERO (Fig. 1). We supported the grant application for the I3D:bio project in 2020/2021 by serving as a use case for a community-oriented implementation of OMERO. Within this project, we started our internal administrative preparations in parallel to the start of the I3D:bio project’s funding phase. In addition, we decided to initiate the implementation of OMERO in our facility by coupling our RDM management initiative to the introduction of a new imaging instrument (i.e., to an automated slide scanner). This opened the exciting opportunity to both facilitate a seamless adoption of new practices by our users and test the handling of large and complex datasets. Administrative preparations during this initiation phase included planning a small pilot project with respect to the temporal, financial, personnel, and infrastructural conditions available at this time.

**Fig. 1:**
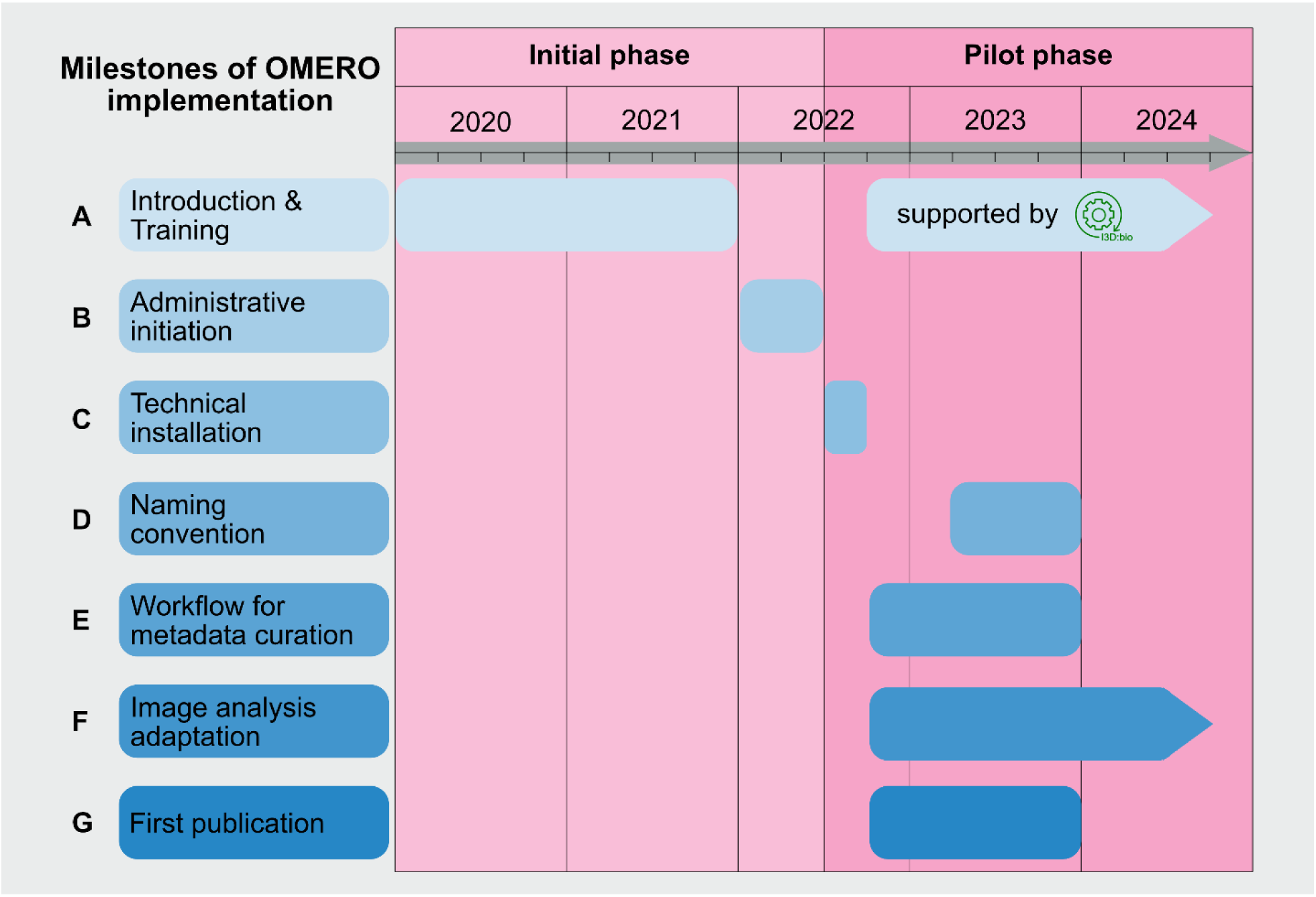
Milestones of OMERO implementation. Illustration of the overall process of implementing OMERO at the CFCI during the initiation phase, with the focus on the administrative organization and the pilot phase with the active usage of OMERO within different use cases. (**A**) Introduction and training covered two different periods: first, initial awareness and general education in the field of RDM and OMERO; and second, active practical training and learning after the installation of OMERO as actively supported by the I3D:bio team. (**B**) Six-month phase of administrative processes. This was an important milestone as resource allocation enabled the start of the technical installation of the OMERO instance. (**C**) Completed over a period of 12 weeks challenges like accessibility of the OMERO instance within the medical domain and beyond as well as different user authentications were covered. Due to the group’s lack of expertise in both RDM in general and OMERO specifically, multiple processes began in parallel. (**D**) Nine-month development phase of a research group-specific naming convention. This phase was necessary for closely linking the newly acquired skills to the in parallel-developed workflows for metadata curation. (**E**) Focused period on understanding the tagging and developing of templates for Key-Value-pair annotations. (**F**) Parallel integration of user-specific data analysis processes from the beginning. This very time-consuming process required specialized knowledge in image analysis and Python programming. This phase is still ongoing. (**G**) RDM of the data related to the PERIKLES project. Following a successful training and annotation phase, this project as part of the four pilot studies was the first one at the medical campus with figures created by OMERO.figure for a scientific publication that is now published.^8^

The conception also required a close cooperation with the IT department from the start to ensure the necessary technical support and to allocate person-time for the installation and maintenance of the new system. This six-month bottom-up initiation phase involved multiple meetings and was a prerequisite for a coordinated strategic decision to implement OMERO. Consequently, we had established an ideal scalable environment to begin the two-year pilot phase in 2022 involving four initial sub-projects.

## Pilot projects

Within the course of implementing OMERO, four projects on campus were chosen reflecting the complexity in multi-dimensional microscopy and the diversity of other types of processed data within our user community (Fig. 2). Furthermore, we selected highly motivated users willing to implement a new RDM routine. The first project was a scientific study, in the following described as PERIKLES project, on the biocompatibility of newly designed biomaterials out of pericardial tissue for cardiovascular substitutes.^8^ Therein, a variety of stained histological sections had to be digitally scanned in high resolution and number using the newly acquired slide scanner (Fig. 2A). The requirements and challenges were not only in the scanning of the large data volumes of more than 20 TB but also in handling the complex experiment-specific metadata that had to be linked to each of the numerous images for following analysis. Furthermore, it was challenging to adapt the already established Fiji macros^9^ to the automated image analysis processes for the OMERO framework.^10^

**Fig. 2:**
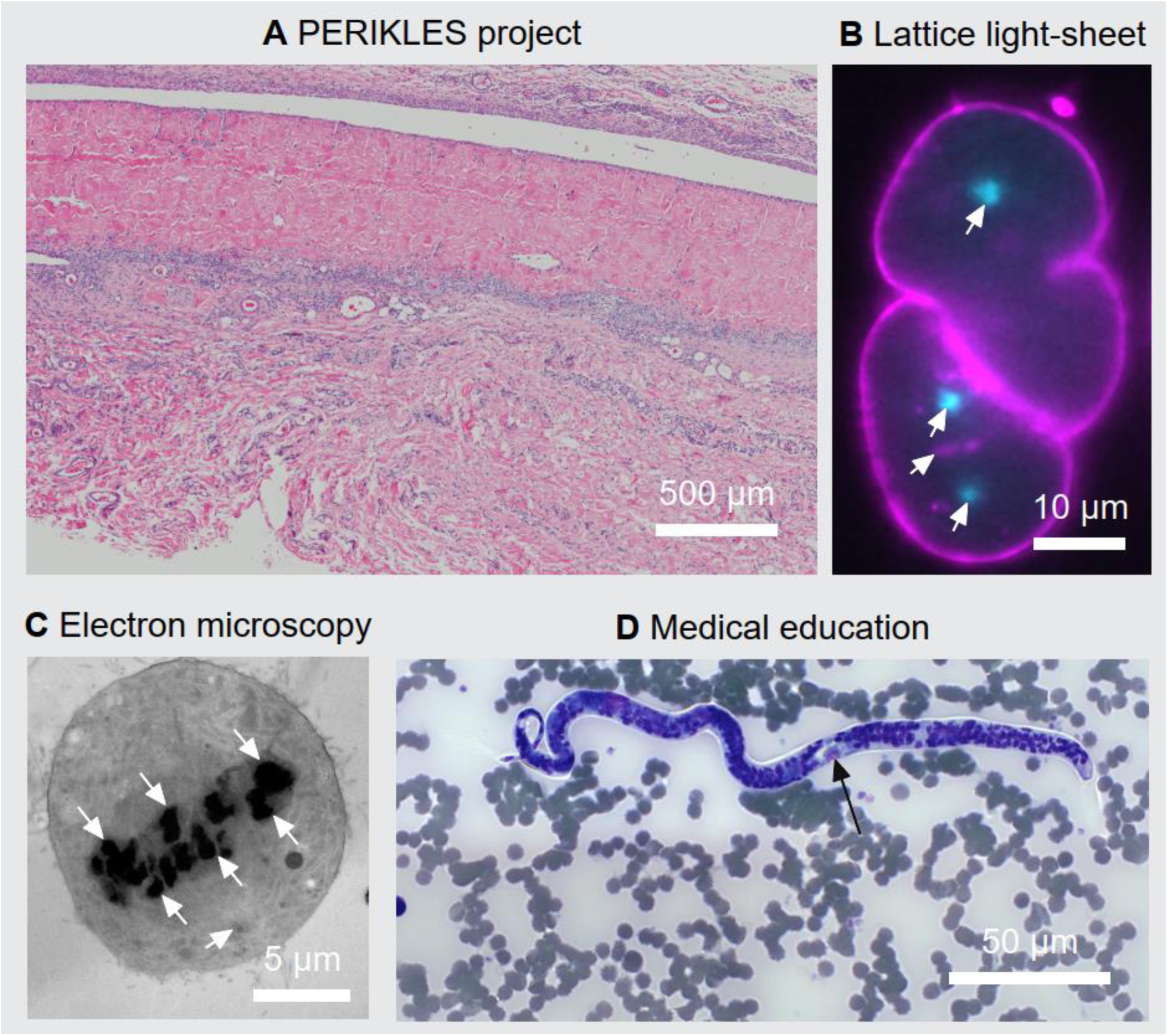
Overview about the pilot projects. This [figure](https://omero.med.tu-dresden.de/figure/file/2425/) was created in OMERO.figure as an example for FAIR-bioimage publication. **A** Light microscopic image of a hematoxylin and eosin stained pericardial implant with granulocyte border area 1 week after subcutaneous implantation into a Sprague Dawley rat. **B** Lattice light-sheet data used to study mitotic spindle scaling in the developing *C. elegans* embryo. The following components are labeled: cell membrane in magenta (PH-domain labeled with mKate2), centrosomes in cyan (gamma-tubulin labeled with GFP, arrows) and histones in magenta (histone H2B labeled with mCherry, arrow head). **C** Transmission electron microscopy image of a RPE-1 cell in metaphase. Chromosomes can be seen in the center of the cell as dark structures (arrows). One centriole pair is visible in the lower part of the cell (arrow head). **D** Light microscopic image of the parasite Loa loa in a blood smear stained with Giemsa staining. The characteristic structure microfilaria is indicated (arrow) and used within lectures and practical courses.

The second pilot project used data obtained by lattice light-sheet microscopy. Here, a small number of experiments visualizing early development in the embryo of the nematode *Caenorhabditis elegans* produced multi-dimensional data including about 3.8 million images and enormous file sizes (approx. 80 GB per experiment). In this case, the challenge was to manage the data in a way that allowed stakeholders to annotate and browse the individual image data with OMERO, while preserving access to the raw data for further processing steps to avoid data duplication as much as possible (Fig. 2B).

In the third project, electron microscopic data of the immortalized human cell line hTERT-RPE-1 was used. Here the ultrastructure of the mitotic spindle was investigated by electron tomography. These data were generated with a number of different electron microscopes resulting in diverse data formats. Due to the complex workflow of data acquisition and image reconstruction this resulted in large file sizes with diverse segmentation results and file formats but at a low sample throughput (Fig. 2C).

Finally, a fourth project was selected from the field of medical education and served as special case for the usage of OMERO in education purposes (Fig. 2D). Here the focus was not on working with large and/or complex data files, but on the parallel, permanent availability of annotated images for up to 300 students per semester in lectures, lab courses and for self-studies (example dataset: omero.med.tu-dresden.de/dataset-1554). An anonymous and unrestricted access of specific images for virtual microscopy and morphological analysis of various microorganisms by the enrolled students had to be embedded into the e-learning platform (OPAL) that is currently used at the Faculty of Medicine at the TU Dresden. In cooperation with the Institute of Medical Microbiology and Virology, digital case studies were selected to be integrated into OMERO.

Throughout the implementation period, we leveraged the support offered by the I3D:bio project. Based on prior work inside the [German BioImaging - Society for Microscopy and Image Analysis](https://gerbi-gmb.de/) (GerBI-GMB), specifically with the working group [Research Data Management for Microscopy](https://gerbi-gmb.de/activities/rdm4mic/) (RDM4mic), members of I3D:bio had established a network of partners with experience in OMERO. This allowed for a coordinated cross-institutional exchange with the I3D:bio team on specific issues where and when they arose. We also used the I3D:bio OMERO training material which was under development, and provided feedback on the basis of our learning experience.^11^

## Getting started with OMERO

Essentially, OMERO is a server to which image data are uploaded and accessed via a set of client applications. The principal application for both browsing the image content and managing the projects is called OMERO.web, a web-based client accessible from any common web browser. Images are most commonly uploaded to the server using the dedicated OMERO.insight desktop client application. In addition, there is a command-line tool, OMERO.cli, which offers a set of advanced functionalities for interacting with the server (e.g., batch import/download/deletion/duplication, custom database queries). OMERO also provides an Application Programming Interface (API) for several programming languages (<u>omero-guides.readthedocs</u>). For the overview and descriptions of used extensions and scripts for OMERO review table 1 in the Supplemental Material.

During the first steps of installing OMERO, we took advantage of the well-described OMERO documentation (openmicroscopy.org/system-requirements) reporting on the minimal technical requirements. The installation of the OMERO environment and the technical support during this initial process were carried out by IT professionals of the Faculty of Medicine. This was done in close collaboration between the CFCI and the Data Integration Center (DIZ) of our faculty. At the beginning of the installation process, a decision had to be made regarding the operating system. In addition to the possibility of a native installation on Linux, there was also the option for an installation via Docker containerization. Because Docker seemed easier to maintain in the long term and offered more flexibility for potential migration or expansion to another system, this model was selected (Fig. 3). The OMERO instance is installed on a virtual machine provided by the DIZ and includes CPU cores, 16 GB of RAM, and initially had two tera-byte (TB) of storage for bioimage data. In line with usage requirements, this storage space has already been increased to the current capacity of 20 TB. With the current total server capacity of 100 TB, the storage can be scaled in line with the usage of the OMERO system. In the future, we will need to expand this capacity even more to a new storage system. However, a migration workflow either using an existing strategy as worked out in the OMERO community or by applying a newly developed solution still has to be discussed.

**Fig. 3:**
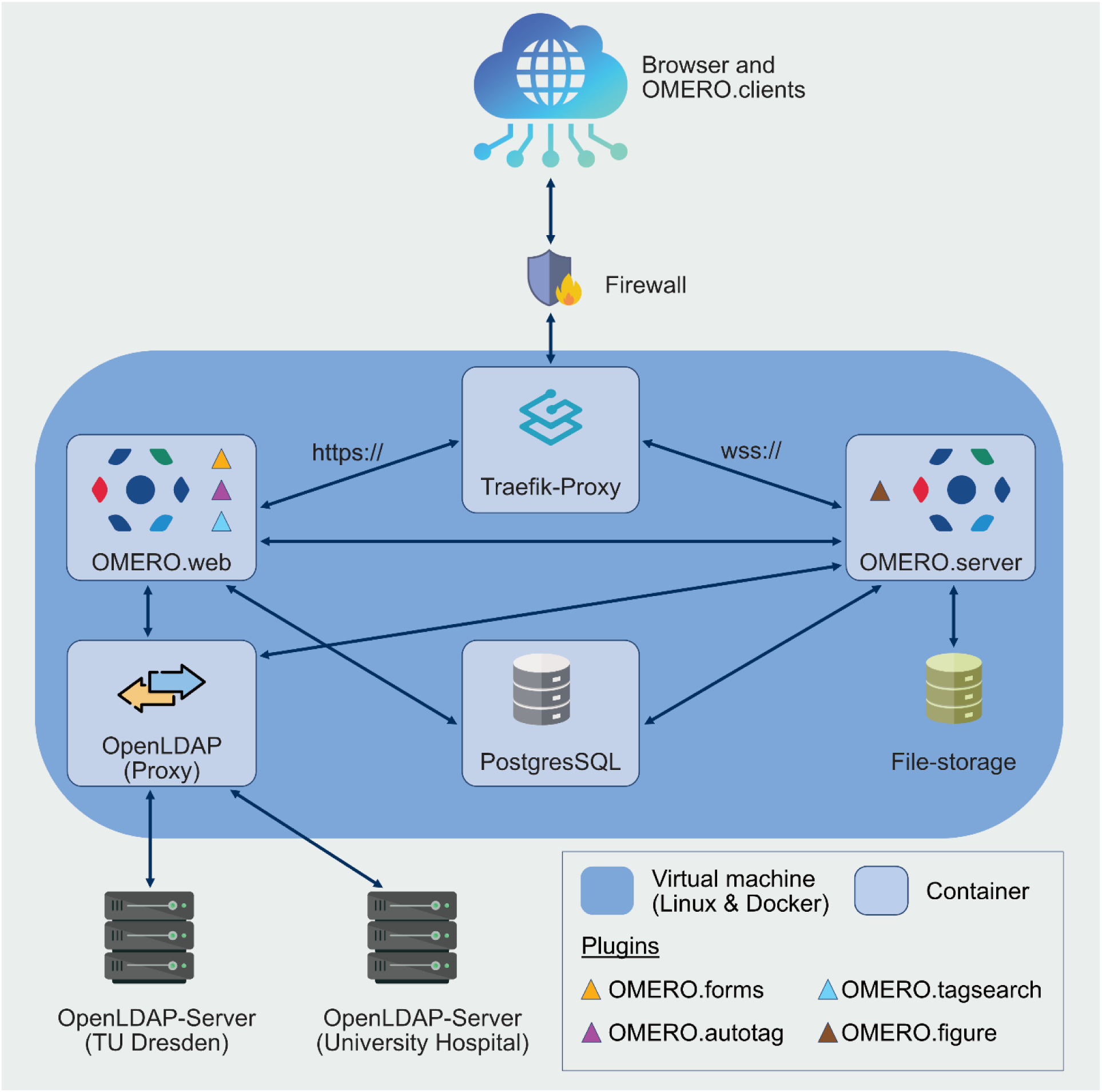
Technical OMERO setup. The Docker stack employed in this installation consisted of both, an OMERO.web (custom docker image) and an OMERO.server container (custom docker image), which are connected to a Traefik-Proxy container (HTTP reverse proxy). Furthermore, the setup is constituted of containers for an OpenLDAP-proxy (OpenLDAP custom image) and a PostgresSQL-database. The OMERO.server container is a custom docker image, based on the image ‘openmicroscopy/omero-server’ (hub.docker.com/omero-server) and has been extended with the OMERO.figure plugin (github.com/omero-figure). The OMERO.web container is also a custom docker image, based on ‘openmicroscopy/omero-web-standalone:5.22’ (hub.docker.com/omero-web-standalone), which has been extended with OMERO.forms (github.com/OMERO.forms), OMERO.autotag (github.com/omero-autotag), and OMERO.tagsearch (https://github.com/German-BioImaging/omero-tagsearch). The user authentication is managed through the custom OPEN-LDAP container. Since an LDAP integration was required for two domains, i.e., for the Faculty of Medicine/University Hospital domain (MED) and the TU Dresden domain (ZIH), the OpenLDAP proxy was necessary to connect both domain trees.

The OMERO.web container, in addition to its use by internal users, is configured for public access to anonymous users. The instructions are available at omero.readthedocs. This configuration allows access to image data and analyses for research team members and collaborators via the plugin OMERO.iviewer. It also allows users to design figures for manuscripts that are entirely created using the OMERO.figure plugin (openmicroscopy.org/omero/figure). The OMERO.web interface has also been set up to allow an automatic login for a ‘PublicUser’, who can access all assigned data, thereby enabling the use of OMERO to make data publicly available. In addition, server settings for open access to published data have been implemented following the instructions in the OMERO sys-admin guide (omero.readthedocs). All settings in the section ‘Configuring public user’ were applied as described there (access date: 06/2024).

### First challenges

After the setup of a new institutional OMERO instance was achieved, a major challenge was the introduction of OMERO in an operating research group with a pre-existing data management structure and associated internal processes. These data management strategies in individual research groups are often not standardized, and they almost always vary across different research teams depending on the specific characteristics and requirements of the different projects. This can present a significant obstacle to the FAIR sharing and usage of the data within further studies.^7^ This issue also impacts the ability of research groups within one institute to comply with the requirement from most funding agencies to store raw and processed data for at least 10 years.

We tackled this problem by developing and commonly agreeing on a group specific naming convention for newly-acquired image data. In our facility and in the associated research group we apply different imaging modalities (light and electron microscopy; Fig. 2B-C). This requires specific information to be included in a valid file name. The development of a naming convention upon which every group member finally agreed was a process that took about nine months of constant discussions, changes, and improvements (see Fig. 1D, Milestones of OMERO implementation at CFCI). As a result, our naming convention now contains a detailed description of how both light and electron microscopic data have to be named in our research group.^12^ These rules are mandatory for file storage either temporarily on file servers or long-term in OMERO. This step of developing a naming convention was certainly necessary to convince group members to ‘break out’ from their own routines and adopt this new naming scheme for a future managing of different and increasingly complex imaging data.

While adoption of the naming convention was a challenge, it was achieved reasonably quickly. A much greater challenge was to organize the data by completely avoiding deep folder/subfolder hierarches. Since OMERO does not provide folders beyond the project/dataset level, users were confronted with the need to understand that data organization in OMERO is maintained by a powerful tagging system. This system is not immediately intuitive for many users, and we observed a tendency to add information to the file names that was originally provided by the folder hierarchy. Such addition of redundant information prevented a proper tagging of individual files and has to be avoided.

To help our researchers adapt to using our new naming convention and the correct tagging of data, we created an example file demonstrating how a project with data and corresponding metadata needs to be tagged in each person’s individual data storage in OMERO ([Light microscopy data](https://omero.med.tu-dresden.de/webclient/?show=dataset-1552), [Electron microscopy data](https://omero.med.tu-dresden.de/webclient/?show=dataset-1555)). For example, in the large-scale PERIKLES project, researchers typically need to work with ID lists containing animal ID, gender and several other experimental conditions. These complex lists can be replaced by Key-Value pairs and tags (organized in tag sets). At the beginning of a new project, one has to create tags covering all the information as usually given in the ‘traditional’ ID lists (see section ‘metadata curation’). When we started to annotate our data within the PERIKLES project, the use of tag sets combining multiple tags was very helpful. In consequence, once a researcher is familiar with the OMERO.web two-level hierarchy, data can be structured and displayed conveniently using the user-friendly tagging system, thus replacing the need for additional and often very complex lists in Excel spreadsheets.

## Workflow development

### Metadata curation

As previously mentioned, the significant conceptual leap for OMERO users was to understand that proper data organization is achieved through tags and tag sets rather than a folder/subfolder structure. The key question in the workflow was when tagging should be performed. Currently, we aim to upload data from the microscope computer or after file creation and then apply the respective tags as soon as possible. This requires a certain level of discipline among group members but is necessary to make data searchable and findable in the future. In projects where many scans were performed in a short period, like the PERIKLES project, tagging directly during upload using the ‘file naming & tags’ function in OMERO.insight proved to be very efficient. In addition, we require all users to provide a short description for each tag, to ensure clarity and avoid confusion.

For existing data with various and individual folder and subfolder structures, trainers need to provide assistance in structuring and annotating them with the perspective to upload data from current projects to OMERO. Here current projects can serve as examples of good practice in RDM for other users at the same institute, especially if annotation templates and file-naming conventions are established during the annotation process. If there are already project- or even technology-specific naming conventions within the research group, these can be adapted through automatic batch annotation using tags derived from the file path from which the images were imported. The OMERO.autotag plugin (https://github.com/German-BioImaging/omero-autotag) reads these paths and subsequently provides the user with a series of tags. We greatly appreciate this function because it facilitates a smooth transition from local file system data management to an OMERO-based data management. Furthermore, handling data in OMERO can be improved by using the OMERO.tagsearch plugin (github.com/omero-tagsearch), which allows searching for tag annotations directly from the OMERO.web interface.

Another feature that aids data reproducibility is the use of Key-Value pairs to document experimental details that lead to the respective microscope image. To annotate image data with Key-Value pairs, spreadsheets can be used and several templates like the REMBI template^13^ are available to ensure compliance with the FAIR principles. Since from our experience it was inconvenient to use those complex templates, as many keys were not relevant to the given project or imaging platform, users should be assisted here. We tried to filter the multiple fields to an essential set of keys to provide values for the experimental setup in the respective projects.^14–16^ The annotations from these templates can be easily imported into OMERO, enabling users to annotate their datasets with metadata in bulk. This is achieved through the annotation scripts (github.com/omero/annotation_scripts/) in OMERO.web, which provides an interface to supply script parameters. Although this approach reduces the workflow complexity for users with limited computational expertise, we have noticed here typical handling errors as listed in the Supplemental Material (Supplemental Material Practical usage experiences).

The ability to add hyperlinks to the Key-Value table offers a flexible way to link electronic lab notebook (ELN) pages directly to the image data. Hyperlinks can also enhance data findability by enriching Key-Value pair metadata with terms and links to the relevant ontologies. For instance, ontology terms for imaging methods, biological entities and organisms were employed for the PERIKLES project with the following links obolibrary.org/obo/FBbi_00000243, obolibrary.org/obo/NCIT_C13005 and obolibrary.org/obo/NCBITaxon_27592, respectively. The Ontobee database was employed to find appropriate ontologies and terms for each annotation.^17^ Moreover, microscope-specific metadata, such as focus strategies or tissue detection settings at the slide scanner system, were also added as Key-Value pairs^15^ to include experimental parameters that are not included in the raw image metadata.

The annotation process described above allows users to establish a robust data management workflow, thus making it easier for the users to implement new data management plans for their imaging studies. Additionally, data curation by tagging and Key-Value pair annotation is very user-friendly, allowing for continued adoption of data annotation practices. The ease of use enabled our imaging facility personnel to efficiently train new users in applying these processes to their data. In perspective, the design of these annotation workflows and templates could greatly benefit from the assistance of dedicated data stewards experienced in RDM for imaging data. The data steward’s expertise and advice could further streamline adopting workflows for existing projects and ensure their inclusion in data management plans for new projects. In the future, this needs to be implemented in our imaging facility.

### Data sharing for collaborations

It is very important to consider the different levels of access to image data within a research group. In addition, the sharing of unpublished data with internal and/or external collaborators must be considered in the data management plan for both pre-existing, current and new projects. This is necessary not only to ensure data integrity and confidentiality but also to facilitate efficient communication between collaborators across project workflows. The initial step in planning data sharing in OMERO is to determine the access levels required by internal team members. This is essential to ensure that only the required users can access, annotate or delete the data. These permissions are set at the group level in OMERO, meaning all group members, except for the group owner and the owner of the data, share the same permissions. Since both users and access requirements can vary between projects, it is recommended to use a separate OMERO group for each collaboration project rather than relying on a single OMERO group for each research group.

Access for collaborators from other research groups, whether within or outside the institute, can be restricted only to the datasets required for the collaboration. The sharing of data with collaborators from the same institute can be managed by the group owners, either independently or with assistance from trainers or the OMERO administrator at the institute. This assistance might be needed, if the user is not yet proficient with the OMERO.web interface, a new group needs to be created, or a new user account needs to be added. However, this last task can be fully automated if the OMERO instance is set up to query the institute’s LDAP server for authentication. This greatly reduces the workload for administrators. Data sharing is generally more complex with collaborators from different institutions, as access to the institute network might be required. There are several possible solutions for this matter. For instance, institutional ‘guest’ accounts to the collaborators might be issued, providing access through Virtual Private Networks (VPNs). Alternatively, the OMERO instance might be accessible from outside by the institute’s internal network. Each of these solutions has different IT requirements and security implications, thus they require extensive planning and collaboration with the institute’s IT department, ideally before installation of the OMERO instance.

Another crucial aspect to consider when planning data sharing is how to avoid data duplication. While the cases discussed above can be addressed by creating new groups in OMERO and moving the data to the new groups, data that are required for more than one project cannot be managed in this way. This is because the projects might involve other datasets with different access requirements, which means that the data would need to be located in more than one OMERO group, potentially leading to data duplication. Data duplication can be avoided by creation of links. This can be achieved by using either the CLI tool omero-cli-duplicate (pypi.org/omero-cli-duplicate) or by reimporting the images a second time using symlinks from the image stored in the OMERO ManagedRepository (omero.readthedocs). In practice, only a new database entry is created, which is linked to the same image file on the server. This “virtual” duplication ensures an independent possibility to work with the image in OMERO separately (e.g., by tagging, adding Key-Value pairs or other kinds of annotations). These independent entries can then be transferred to the group of a collaborator so that the data can be shared efficiently. The downside of these solutions is that they require the user to be proficient with the command line. Administrative assistance might also be required.

### Image Processing

Introducing OMERO into ongoing projects, i.e., when data have already been generated and analysis workflows been established, can come with several challenges. These challenges include constraints on the choice of the programming language and the image analysis software. However, it is important to minimize changes to the original workflow when adapting it to OMERO. This ensures consistency within the project itself and makes the transition as smooth as possible for the users. A prerequisite for the introduction of OMERO to both pre-existing and new workflows is its interoperability with various imaging analysis software packages. As previously described, OMERO is primarily designed for data management and metadata curation, providing limited image processing functionalities out of the box. This limitation is fully compensated by the availability of APIs and plugins for the most common programming languages and image processing software, including Fiji^18^, napari^19^, QuPath^20^ and Cell Profiler.^21^

Effective communication between trainer and user is crucial to ensure that all requirements of the original workflow are identified and implemented in the scripts, tools and/or protocols to be applied after the introduction of OMERO. It is important to frequently test the workflow in close collaboration with the users to gather necessary feedback. For instance, one issue we have encountered with many image analysis plugins for OMERO is the inability to open large images that do not fit in the memory available on a typical workstation or even in a dedicated image-analysis desktop PC. While some software, like QuPath, can open whole-slide images by leveraging multi-resolution pyramidal file formats, other project requirements might prevent the use of such tools. This is particularly true for ongoing projects where data analysis has already been partially performed or where analysis scripts and workflows have already been developed. To enable one of these workflows in Fiji, we adapted a pre-existing macro from the OME team (github.com/open_image_after_download.groovy). This macro downloads an image from OMERO and temporarily saves it to the local hard drive before opening it. We modified it to allow opening of the image at a lower level of resolution and account for a different pixel size in the downstream analysis (github.com/FixSizeDownloader.groovy).

Another example of an adapted workflow is the PERIKLES use case. For this project, we modified a Fiji macro^9^ for counting cell nuclei on hematoxylin and eosin (HE)-stained slide scanner images (github.com/CountCellsOMERO.ijm). This macro was adapted using the OMERO Macro Extensions plugin (github.com/omero_macro-extensions)^10^ enabling the macro to leverage tags and regions of interest (ROIs) created through the OMERO.web interface. This allowed for selecting of the images to process and limited the areas to be processed within each image. The results of the analysis were then imported to OMERO as cell ROIs and measurement tables linked to the relevant images, and made them accessible for viewing and available for download and further analyses (Fig. 4).

**Fig. 4:**
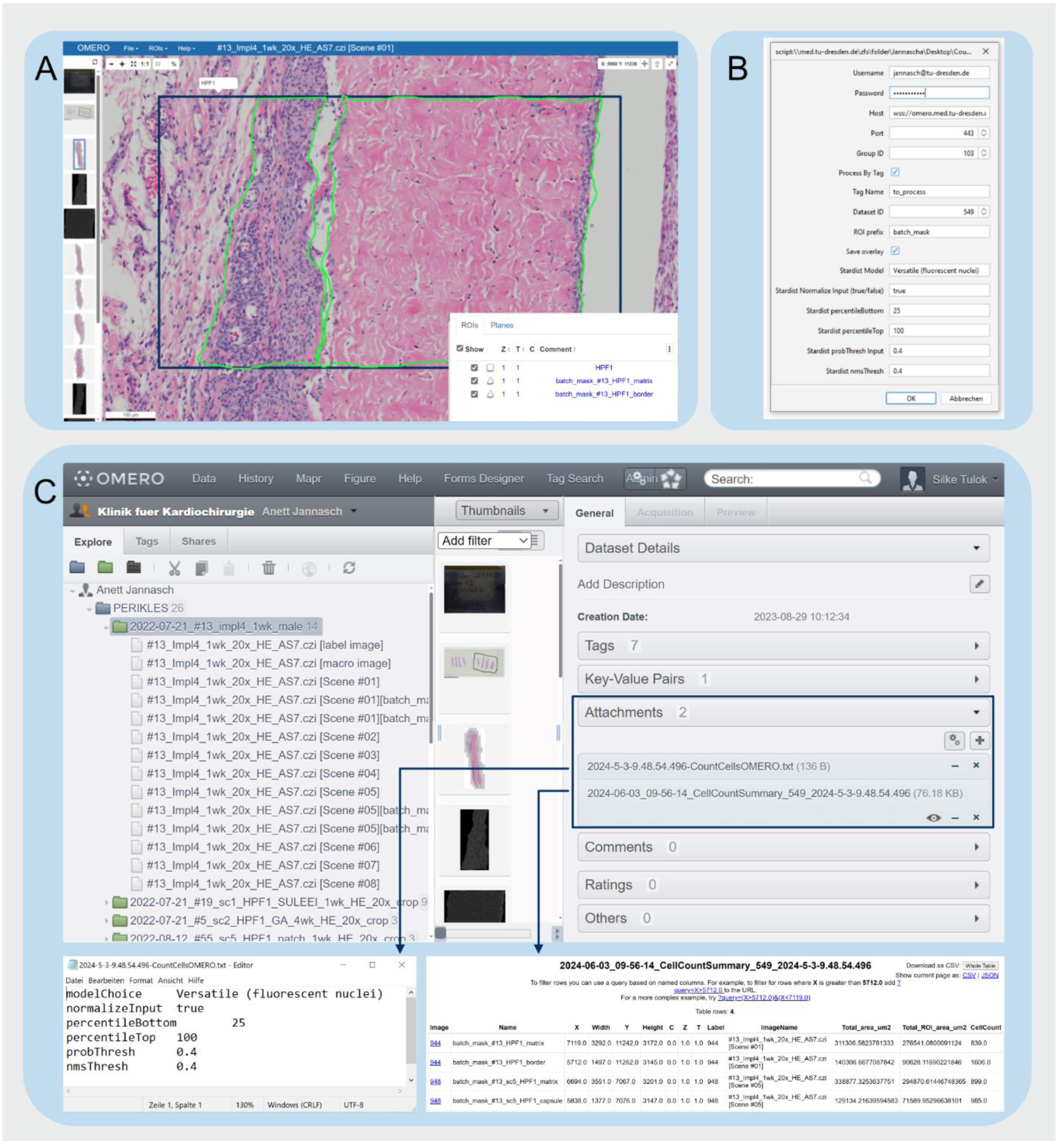
Workflow for the ‘CountCellsOMERO’ macro. (**A**) Screenshot of the OMERO.web interface default viewer showing a representative image and the ROIs employed to select the area for image processing. For the displayed PERIKLES study, it was necessary to have an outline rectangle (dark blue) to ensure the mandatory size of the high-power field (HPF; 1000 x 500 µm^2^ according to ISO 10993-6:2007^22^). Here, one aspect of the project was to quantify cells within the fibrous capsule and implant matrix. Further segmentation as polygons (in green) within the rectangles were done to define both structures. (**B**) Screenshot of the Fiji dialog window shown at the beginning of the macro execution for parameter input. OMERO Login credentials, group and dataset ID needed to be selected. To choose polygons and not rectangles to be processed, we added the prefix term ‘batch_mask…’ and inserted this term in the dialog window. To process images of multiple datasets in a batch, we tagged all favored images with ‘to process’. (**C**) Screenshot of the OMERO.web interface showing the parameters and measurements from the macro saved as an attachment in OMERO. Parameters and measurements are displayed as a table (bottom left and right, respectively).

A drawback of this approach is the limited capability of OMERO to manage and visualize large numbers of ROIs, which becomes impractical above a few hundred ROIs. A workaround here is the use of segmentation masks, which can define several thousand regions for each image, although they are currently not supported very well for visualization in OMERO. This issue will likely be resolved by the adoption of next-generation file formats such as OME-NGFF.^23^ This will enable the storage of these masks together with the original image in a file highly optimized for direct access by a large number of users from cloud or object storage.

Finally, comprehensive documentation should be compiled for each step of the workflow and ideally employed by the user when carrying out testing of the workflow. This will allow users and trainers to identify steps and tools which need further adjustments or clarifications, thus enabling users to independently carry out the new workflow from the very beginning. This documentation will ensure that specific steps of the developed workflow can be re-used and reproduced by future users after the project has ended. If the new workflow should require the development of new scripts and tools, a version control system should be employed to keep track of the script versions used and any subsequent modifications that have been developed. The new code can then be made available either internally or externally on DevOps platforms like GitHub (https://github.com) or GitLab (https://about.gitlab.com). To streamline application by new users, the documentation for the workflow could also be hosted on the platform along with the scripts. An example of such a repository, containing Fiji macros developed for the PERIKLES study, can be found at: github.com/fiji_omero_workflows.

### Publishing

An important feature of OMERO is the ability to provide datasets and figures for publication in scientific journals. Many research groups have agreed on the use of standard graphic software. Importantly, information about the metadata behind composite figures is usually known only by the group members who created the figures for a specific publication. For the publication of the results of the PERIKLES study^8^, OMERO allowed us to make the corresponding histology data available to both the reviewers and the readers. This was appreciated by the journal, APL Materials. Briefly, we would like to share our workflow of publishing data by using OMERO.

We used the application called OMERO.figure, built into the OMERO.client, to create the respective figures for publication. This was very convenient as images were directly used with the correct scaling in OMERO, eliminating the need for duplication on local computers to create high-resolution composite figures. The handling of this application is easy and intuitive, as helpful tutorials are available (openmicroscopy.org/omero/figure). These benefits are also strong arguments to convince other colleagues and researchers to get started with OMERO. We strongly recommend to start with importing image data obtained within current projects and there especially with data intended to be published. From our perspective, this is an ideal starting point for the use of OMERO and encourages FAIR data publication practices within the research groups. The tagging feature was particularly useful here, as labels were generated directly from the given tags. The obtained figures were then saved as .pdf, .tiff files or as a new OMERO image. The link to the figure and a list of the containing images with their individual OMERO links were initially attached to the datasets within the own group. Next, a ‘PublicUser’ group was created (see section Getting started with OMERO) with the ‘PublicUser’ as a group member and read-only permissions. Subsequently, the figures and the corresponding datasets were duplicated, creating a new entry in the SQL-database that was linked to the original image file (see section Data sharing for collaborations). The duplicated datasets were then moved to the ‘PublicUser’ group, and the dataset owner was changed to the ‘PublicUser’. Finally, as the image IDs within the duplicated datasets had changed, they needed to be corrected in the public figure (Fig. 2). From here on, the new figure link could be provided for the manuscript.

For the implementation of OMERO in the microbiology teaching to students enrolled in medicine, digital images of stained slides and solid medium plates of various microbial samples (e.g., bacteria, fungi and parasites) were used. After acquisition of microscopic images, they were stored and properly annotated with relevant information in OMERO. As these images were intended to be shared within a group of students without giving them permissions as OMERO users, the data needed to be provided publicly. Again, a new ‘Public User’ group was created and the data was duplicated virtually there as described above. To enhance the e-learning aspect, it was possible, together with the local faculty’s student e-learning platform ‘OPAL’, to provide the students access to specific images through hyperlinks, embedded into individual chapters of the course. By virtual microscopic analysis, the students were asked to identify specific microorganisms in the given samples, thus linking causative pathogens to selected case studies. For this purpose, non-annotated images of given specific case descriptions on the e-learning platform were used in synchronous virtual microscopy courses. After the end of the course, a fully annotated version of the images was provided (again by data duplication), encouraging independent self-studies by the participating students.

## Discussion

Despite many advantages and benefits for both facility staff and users, the implementation of RDM still represents a huge hurdle for most researchers. This particularly concerns small laboratories with low capacities and/or expertise to sustain resources and introduce workflows for FAIR data management.^24^ Hence, there is an urgent need for support from the local IT department as well as from data stewards. This is true not only for the installation and setup but also for the maintenance of the RDM infrastructure and accompanying user training.^25^

This need for support was the driving force behind the pilot project described here. Our goal was to develop and share a streamlined workflow for RDM that could alleviate the burden on researchers as mentioned above and facilitate better data management practices in bioimaging facilities. From the beginning, it was our intention to start with a well-described and supported system, OMERO, for bioimage data management. To a large extent, we benefited here from the experiences of the microscopy community in introducing and using OMERO.^2,5,26,27^ The recently published comprehensive facility perspective on adopting the FAIR principles provides high-level recommendations on implementing a bioimaging RDM system.^2^ Here, we present the perspective from a dedicated user case. Our article describes in detail the bottom-up process on selected pilot projects, including the implementation of OMERO in diverse research areas and the teaching activities at our research campus. In, addition, we also shared developed protocols. As we started the project on a small level with the option to expand and upscale the use of OMERO, the parallel community support including functionality development for OMERO was crucial for getting started with a professional RDM in the CFCI.^10,28,29^

The strategy for implementing OMERO in the context of this pilot study was initiated as a bottom-up process by the CFCI. Our initiative then created further awareness at the faculty level for the importance of a management strategy for the handling of bioimaging data. At this point, the need for additional personnel and technical resources has not been finally discussed. Specifically, aspects of RDM ranging from data analysis, sharing to publication are currently under discussion to identify solutions within the scope of our pilot studies. This initial phase is crucial for the subsequent and targeted institutionalization of OMERO as a central bioimaging data management system. The success in improving data handling and alleviating workload for RDM experienced by pilot users seems to stimulate the uptake of OMERO into the routines of other groups within the collaborative environment of a research institute. Our long-term aim is to convert this initiative into a top-down approach, thus enabling the recruitment of additional personnel to offer faculty-wide technical support. Such a central top-down process will be crucial to motivate a larger group of scientists in research groups to change their old lab routines and develop new and future-proven data management workflows.

A notable example of successful RDM implementation is the PERIKLES project. This research project was the first in its domain at the campus of the Faculty of Medicine in Dresden, which successfully incorporated the OMERO platform into its publication process. OMERO enabled the authors to share annotated histological images with team members and collaborators, thus shaping an effective evaluation of biocompatibility for a new candidate biomaterial for cardiovascular substitutes in a transparent and comprehensible way. This transparency is crucial for data reproducibility and the peer review process allowing colleagues in the field to verify and build upon the findings. Moreover, the open data policy during the PERIKLES project was recognized by the editorial office of APL Materials. This appreciation underscores the value of open data practices in enhancing the credibility and impact of scientific research.

Furthermore, we successfully used the newly installed OMERO platform for teaching students enrolled in medicine. Due to its public accessibility and implementation in the local student e-learning platform OPAL, students have gained great benefit from using annotated imaging examples for lectures, practical sessions, and exam preparations. This newly established virtual microscopy library will be increasingly used for other topics to be taught as well as for an internal training of employees, researchers and also for student helpers.

The success of this initiative relied on a close collaboration with the I3D:bio project, which supported the deployment of the OMERO server. This collaboration served as a use case for I3D:bio, aiming: first, to develop training material and guidelines for a core facility, and second to demonstrate the deployment of OMERO in a microscopy facility with no prior experience in this RDM approach. Since the I3D:bio project began concurrently with this collaboration, it was a learning experience for both parties. On the one hand, the CFCI staff gained technical knowledge in the operation of OMERO and in implementing data management concepts from dedicated training sessions with the I3D:bio team. On the other hand, the I3D:bio’s teaching evolved as their expertise with OMERO grew and benefited from the feedback it received from the CFCI. This led I3D:bio to publish their training material^11^ and develop further open-source projects that were previously used for the CFCI projects (omero-scripts CSV, simple_omero-macro).

Projects like I3D:bio will continue to develop training materials and conduct workshops on a larger scale to train core facility personnel in implementing and using OMERO. Unfortunately, I3D:bio will not have the resources in the future to support additional facilities as intensively as it was done with the CFCI and its pilot projects. Therefore, it is crucial here to emphasize the urgent need for dedicated personnel specializing in a data management at core facilities. These individuals will need to take over the task of supporting local projects with data management, while remaining connected to the broader data management community for further advising, training, and sharing their expertise. Besides support through I3D:bio, the currently developing National Research Data Infrastructure (NFDI) provides resources and data stewardship to support researchers and core facilities in bioimaging research data management (nfdi4bioimage.de/help-desk). To upscale the successful implementation of OMERO, the CFCI imaging facility could serve as an interface for building a data management system following the FAIR principles on the entire campus of the Faculty of Medicine of the TU Dresden. With the successful initiation of OMERO for slide scanner data, many users already benefited from our developed templates^15^ and Python scripts. This approach will be further developed for other microscopy systems and techniques to make it as easy as possible for future users to smoothly adopt OMERO with maximum functionality.

In summary, the CFCI is ideally positioned for the successful implementation of this pilot project due to its structural integration within the Faculty of Medicine, its personnel expertise in the field of light and electron microscopy, and its existing network structures. To achieve the necessary subsequent faculty-wide scaling to a multi-user environment, it is essential to secure personnel infrastructure and the associated IT resources. This must be aligned with central faculty decisions in the field of research data management.

## Conclusions

Implementing a professional RDM system presents challenges, primarily due to the time and expertise required from researchers and core facility staff. Local support by imaging facilities and IT specialists, and tailored solutions are crucial for the effective integration of RDM into research workflows, thus ultimately contributing to the advancement of open science and data-driven discoveries. With our experiences and procedures during the whole process ranging from administrative issues, including installation processes, workflow developments and image processing up to publication, other local and national bioimaging communities will certainly benefit by setting up their own OMERO instance.

## Acknowledgments

We thank Dr. Vanessa Fuchs from the NFDI4BioImage for her great support during the development of data annotation templates. We acknowledge the advice of Dr. Robert Haase from the NFDI4Bioimage within the startup and initiation process of the project. We thank all members of the AG Müller-Reichert (Core Facility Cellular Imaging CFCI) for the fruitful discussions and beneficial contributions. We would like to thank Maria Feilmeier, Jennifer Mittag and Dominic Salminger from the research department of cardiac surgery for their excellent technical assistance with the histological analysis in the PERIKLES project. We acknowledge the allocation of IT infrastructures and personnel resources as well as the administrative support from Prof. Martin Sedlmayr (CIO of the Center for Medical Informatics of the University Medical Center Dresden). Furthermore, we thank Dr. Peter Dieterich (Head of IT Department of the Faculty of Medicine Dresden) for initiating the collaboration between CFCI and the IT department of the Faculty of Medicine. We acknowledge the exchange with local and national collaborative networks in particular [BioDIP](http://www.biodip.de/) and GerBI-GMB helping to develop and shape the project. Research in the Müller-Reichert lab is supported by grants from the Deutsche Forschungsgemeinschaft (DFG; MU1423/8-3, grant no. 258577783 and MU1423/10-3, grant no. 282354882). The Müller-Reichert lab also acknowledges support from the CCBX program of the Center for Computational Biology of the Flatiron Institute (NY, USA). The PERIKLES project was funded by the European Regional Development Fund (EFRE) and the Freistaat Sachsen (grant no. 100367226). The contributions from Dr. Tom Boissonnet and Dr. Christian Schmidt are funded by the Deutsche Forschungsgemeinschaft (DFG, German Research Foundation), project I3D:bio – grant no. 462231789. Dr. Michele Bortolomeazzi is funded by the Deutsche Forschungsgemeinschaft (DFG) under the National Research Data Infrastructure - NFDI 46/1 – grant no. 501864659.

