## Supplemental Material for "Setting up an institutional OMERO environment for bioimage data: perspectives from both facility staff and users"

### Supplemental Materials

Table 1: Extensions, customizations and scripts for OMERO used at Core Facility Cellular Imaging (modified after Zobel et al.[1])

| **Name** | **Type** | **Description** | **Source** |
|---|---|---|---|
| OMERO.web | Client based on Python 3 | Provides a web-based user interface and JSON Application Programming Interface (API) | omero.readthedocs.io |
| OMERO.insight | Desktop client application | Graphical desktop client application with a user-friendly interface to interact with the OMERO.server (upload, tagging, …) | omero-guides.readthedocs.io/import-desktop- |
| OMERO.iviewer | OMERO.web app | Viewer with bioimage visualization, annotation and analysis functionalities | openmicroscopy.org/iviewer |
| OMERO.figure | OMERO.web app | Creation and management of figures from multi-dimensional images stored in OMERO | openmicroscopy.org/figure |
| OMERO.forms | Extension to OMERO.web | To enhance metadata input and provide provenance, which allows users to create, edit, and assign forms for use within specific groups | github.com/OMERO.forms |

| | | | |
|---|---|---|---|
| OMERO.autotag | Plugin | Automatically tagging of images or datasets based on their metadata or image content | github.com/omero-autotag |
| OMERO.tagsearch | Plugin | Enhances the ability to search and filter images, datasets, and projects using tags | github.com/omero-tagsearch |
| OMERO.client | Client (command-line based) | Programmatic client library that to interacts with the OMERO.server using various programming languages, such as Python, Java, and MATLAB | docs.openmicroscopy.org/cli |
| OMERO.cli | Command-line plugin | Command-line tool | omero.readthedocs.io/cli |
| OMERO-cli-duplicate | Command-line plugin | Duplication of objects in OMERO, without duplicating image data | github.com/omero-cli-duplicate |
| OMERO Macro Extensions | Plugin & Scripts | enables to leverage tags and regions of interest (ROIs) created through the OMERO.web interface | doi.org/10.12688/f1000research.110385.2 |
| Download groovy script | Script for Fiji | Fiji macro that downloads an image from OMERO and temporarily saves it to the local hard drive before opening it in Fiji | github.com/open_image_after_download.goovy |
| FixSizeDownloader macro | Script for Fiji | Modified Fiji script to allow opening of image at lower resolution level (image pyramid) and account for different pixel size | github.com/FixSizeDownloader.groovy |

| | | | |
|---|---|---|---|
| CountCellsOMERO macro | Script for Fiji | Modified Fiji macro for counting cell nuclei on hematoxylin and eosin stained slide scanner images | github.com/CountCellsOMERO.ijm |

1 Zobel, T., Weischer, S. & Wendt, J. (2022) OMERO for microscopy reserch data management-A use case example from the Münster Imaging Network. Wiley Analytical Science News https://analyticalscience.wiley.com/content/article-do/omero-microscopy-research-data-management

### Practical usage experiences

In this section, we summarized some more practical usage experiences, which might be interesting and useful by starting an OMERO environment.

#### *Data upload of "big datasets"*

We encountered problems during the data upload to of our Lattice-light sheet data to OMERO. The upload of that dataset (80GB) caused in a timeout in the final processing step after the upload procedure, which was caused by the reverse proxys. We could circumvent that by using a different server address with a direct connection between OMERO.web and OMERO.server. Thereby all other proxy containers were avoided. The disadvantage of that procedure is, that the data upload was only possible within our medical domain.

#### *Orphaned files after failed upload*

We also discovered after a failed upload, that there are still files saved on OMERO.server in the user folder. Those files are not visible in OMERO.web but still block space on the entire storage. The problem is, only after a successful upload the file gets a database entry with an image ID and is visible via OMERO.web. Orphaned files are still on the server but "invisible" to the users.
Over time every failed upload might accumulate a large data volume that is unusable and undetectable via OMERO.web. With help of the community, we found a way to detect those files on the server (https://forum.image.sc/t/orphaned-files-on-the-omero-server-after-failed-upload/97184) via the command line. This list of files can also be deleted afterwards via deletion commands. Here, we will work in future on an automated solution to clean up server space, which can be activated from time to time by the responsible data stewards or IT support.

#### *Tagging*

Tags always belong to the person, who created them. That means, if the owner of a project, dataset, and/or image is changed to another group all tags will be lost. This was important for us during our publishing and sharing workflows. Therefore, we had to re-tag the public data. From our perspective, it would be very helpful to have different tagging categories, which can be transferred in-between different groups and owners (e.g. global tags, group-specific tags, and user-specific tags).

#### *Key-Value Pairs*

For the metadata annotation with Key-Value pairs the simplest method was creating a csv-template to upload via an OMERO annotation script (https://github.com/ome/omero-scripts/tree/develop/omero/annotation_scripts). In developing our workflow, we discovered a problem: the created csv-files were not able to populate the metadata. The solution to the problem was the 8-Bit UCS Transformation Format (UTF-8)-Coding. We discovered that some Excel-versions (in our case an older version of Microsoft Office) don't save csv-files in that code. If such files have the wrong code, some characters might be not able to decode and the automated annotation via this annotation script will not run. Detailed descriptions can be found here (10.5281/zenodo.12547566, 10.5281/zenodo.12578084, 10.5281/zenodo.12546808).

##### *Batch processing by using tags*

One common pitfall we experienced was to forget to set the ‚to_process' tag before or after the macro ‚CountCellsOMERO' was executed. In the first case one simple does not get the analysis on the images and in the second case, a mass of ROIs will be detected again and implemented in that image. To delete big amounts of unwanted ROIs easily, open OMERO.insight client, select the image and choose 'ROI Tool…' in the 'Controls' tab. Same holds true for tables, go to OMERO.insight client, select image and delete attachments.

##### *Sharing data with collaborators*

If large data are intended to be shared with collaborators and they need to download the images for further processing steps (e.g. training of deep learning networks). We encountered problems with the download function offered by OMERO.web as the download often stopped before it was completed. A possible workaround we found was to provide the collaborator a temporary guest login and make them member of a collaborator group in which the data set is located. Then the person could login to OMERO.insight, navigate to the data set and start the download this way. We found that the download via the OMERO.insight client was stable enough to complete the download of large files.

##### *Fluorescence Lifetime Imaging Microscopy (FLIM) - Data*

In the moment it's not possible to visualize, and manage FLIM data (in our case acquired with a Leica Stellaris system) to OMERO. This is currently under active development by the OMERO-community. For now, we would not recommend to upload any FLIM data to an OMERO environment, because their deletion is not possible in the OMERO.web and needs to be done via the command line.

##### *Medical Education*

In the field of medical education, we have several ideas for improvements within OMERO. We would recommend the implementation of a selective teaching display mode or settings with limited or restricted user functions. For teaching purposes, it would be useful to direct the user directly to the annotation tab without the need to access other tabs. Ideally, this setting could be set as standard or alternative the other tabs are not displayed. Furthermore, when redirecting users from an external learning platform to a specific image, the user could see and access all other images in the same folder. For a strictly external learning platform driven redirection to images if would be helpful to hide other images in the same OMERO folder from visibility and access by the user.

Another point to improve would be the scaling of image annotations (arrows and text), in particular at low magnifications as such annotations are only partially readable. This could be fixed by setting a certain default zoom level for each individual image.
